# SiGn: An Open-Source Signal-Generating fMRI Phantom for Dynamic Quality Assurance

**DOI:** 10.64898/2026.04.15.717370

**Authors:** Sephora Galea, Brandon C Seychell, Paola Galdi, Thérèse Hunter, Claude J Bajada

## Abstract

Functional magnetic resonance imaging (fMRI) quality assurance has traditionally relied on static, geometrically regular phantoms that cannot generate the dynamic signal changes fMRI analysis pipelines are designed to detect. Here we present the Signal Generating (SiGn) anthropomorphic brain phantom, a 3D-printed cortical model derived from an individual participant’s structural MRI, filled with tissue contrast-mimicking agar gels and coupled to a hemin-based infusion system that produces controlled, time-varying 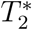-weighted signal changes. We validated the phantom across two scanning sessions on a 3 T Siemens MAGNETOM Vida scanner, demonstrating that hemin infusion produced spatially localised activation detectable by standard general linear model analyses. Because the phantom’s geometry is derived from real participant anatomy, its functional data can be coregistered and spatially normalised to standard brain templates through the same pipeline applied to human data, enabling end-to-end assessment of how each preprocessing step affects a known ground-truth signal. To support adoption and reproducibility, we openly release the full resource, including 3D-printable STL model files, tissue-mimicking gel recipes, the BIDS-formatted dataset, preprocessing and analysis scripts, and a containerised reproducibility workflow; the corresponding archival container image is also deposited on Zenodo. This framework is intended to lower the barrier for other groups to fabricate, scan, and analyse an equivalent device on their own hardware, adapt it to specific research questions, and iteratively improve the design, thereby supporting more rigorous and transparent fMRI quality assurance practices across the neuroimaging community.

## Introduction

Functional magnetic resonance imaging (fMRI) is a cornerstone of human neuroscience, providing non-invasive access to brain function across a large range of applications, from presurgical mapping of eloquent cortex [1–3] to the study of cognition and neuropsychiatric conditions [4–6]. The most widely used fMRI contrast mechanism exploits the blood oxygenation level-dependent (BOLD) effect, in which fast echo-planar imaging (EPI) sequences detect the decrease in paramagnetic deoxyhaemoglobin concentration that accompanies local neural activity via neurovascular coupling [7]. Despite its clinical and scientific utility, the BOLD signal is inherently noisy and susceptible to macrovascular contamination, physiological fluctuations from respiration and cardiac pulsation, and diverse motion-related artefacts [8], all of which impose practical limits on the reliability of fMRI measurements.

Systematic quality assurance (QA) procedures can characterise scanner stability over time, verify that acquisition parameters produce expected image quality, and ensure that preprocessing pipelines perform correctly [9, 10]. Software-based quality control (QC) tools, such as the Automated Processing Quality Control (APQC) system in the AFNI toolbox [9], facilitate structured review of motion estimates, registration, and temporal signal-to-noise ratio, but they operate on data that are already confounded by the physiological variability of the human participant. To obtain a measurement reference that is independent of participant state, the field relies on MRI phantoms: inanimate test objects, standardised or otherwise, that can be scanned repeatedly under controlled conditions to assess scanner performance, calibrate quantitative measurements, and validate new sequences [11–13]. Phantom-based validation of this kind is a well-established methodological standard across medical imaging modalities more broadly, with recent guidance emphasising rigorous design, reporting, and reproducibility as prerequisites for meaningful interpretation [14].

The current benchmark for MRI quality assurance is the NIST/ISMRM system phantom [11], a rigorously characterised, geometrically regular device incorporating calibrated arrays of spheres and fibres to quantify signal-to-noise ratio, geometric distortion, relaxometry accuracy, and slice profile. Alongside it, uniform agar-filled spheres and water cylinders standardised by bodies such as the American College of Radiology (ACR) and the Functional Bioinformatics Research Network (FBIRN) are highly effective for characterising baseline scanner parameters [15, 16]. By providing a homogeneous field environment, these devices ensure that observed fluctuations can be attributed to hardware rather than biology, and their metrological rigour is precisely what makes them the field standard. However, this same geometric simplicity is their limitation for fMRI-specific validation: neither the NIST phantom nor the ACR/FBIRN devices reproduce the tissue boundaries, susceptibility interfaces, and partial-volume effects that characterise real brain data, and none can generate the dynamic signal changes that fMRI analysis pipelines are specifically designed to detect.

More anatomically realistic phantoms have therefore been developed to close this gap. Structured but non-anthropomorphic designs, such as resolution targets or multi-compartment inserts, address some shortcomings of uniform phantoms [11]. Anthropomorphic brain phantoms, typically produced by 3D printing, go further by replicating macroscopic cortical anatomy, including sulci and gyri derived from segmented structural MRI of a real cortical surface, rather than the whole brain volume, providing a test environment that more closely mimics the geometric challenges of a real experiment [17, 18]. Yet even the most anatomically detailed static phantom is of limited use for validating fMRI analysis pipelines, since it produces no dynamic signal against which detection and localisation can be tested.

A significant gap therefore remains: no widely available phantom combines anatomical realism with a controlled, dynamic signal that can serve as ground truth for fMRI pipelines. A small number of “active” phantom designs have been reported. Some use current-carrying conductors to induce local magnetic field perturbations that mimic BOLD-like signal changes [19, 20]; others employ mechanical approaches such as bubble compression to alter 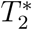 [21]. While these provide adjustable signal modulations, they typically involve cumbersome electrical circuits or produce field perturbations whose spatial profile differs substantially from the distributed susceptibility changes seen in vivo. Dual-compartment gel models have also been developed to represent “active” and “baseline” relaxation states [22], but these tend to use static concentrations of contrast agent and lack the ability to simulate the gradual, temporally varying dilution of paramagnetic species that characterises the haemodynamic response. Furthermore, rigid internal partitions between tissue compartments create signal voids and susceptibility artefacts at boundaries that are not present in biological tissue and that can confound the measurements the phantom is intended to validate [23].

Critically, none of the phantoms described above reproduce the biological mechanism of neurovascular coupling. No current phantom, active or static, can mimic the metabolic and haemodynamic cascade that couples neural activity to changes in blood oxygenation. This is a limitation shared across the entire class of “active” and dynamic phantoms, and it constrains what any such device can claim to validate: phantoms of this kind can test whether acquisition and analysis pipelines correctly detect, localise, and quantify a known, time-locked physical signal change, but they cannot validate the biological specificity or sensitivity of BOLD as a marker of neural activity itself.

The primary aim of this work is to provide the field with an open-source, reproducible platform for generating such controlled, dynamic signal changes within an anatomically realistic phantom, addressing the gap identified above while remaining fully accessible to other groups. Unlike the NIST phantom [11] and other commercial or proprietary anatomical phantoms [17, 18, 23], every component of the design presented here—from the anatomical model and infusion hardware to the tissue-mimicking gel recipes and full analysis pipeline—is released under open licences, allowing other laboratories to reproduce, adapt, and extend it without the cost or access barriers associated with standardised commercial phantoms. Open-source MRI phantom designs have recently begun to emerge, for example, for low-field quantitative imaging [24], demonstrating the value of this approach to reproducibility. However, no open-source phantom to date combines anatomically realistic, brain-shaped geometry with a controllable, dynamic fMRI-relevant signal source.

We refer to this platform as the signal-generating (SiGn) phantom. It combines 3D-printed cortical anatomy, segmented from an individual participant’s structural MRI, with tissue-mimicking agar gels and a hemin-based contrast agent delivered dynamically through an infusion system. Hemin (ferric protoporphyrin IX chloride) produces paramagnetic susceptibility changes that shorten 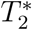 relaxation in a manner qualitatively analogous to the effect of deoxyhaemoglobin in vivo, enabling time-varying signal intensity changes with controllable “on” and “off” states. As an initial demonstration of this platform’s utility, we present a validation study using fMRI acquisition and analysis, in which we test whether standard preprocessing and statistical pipelines correctly detect and localise the time-locked signal changes the phantom produces. This fMRI validation is intended as one demonstration of the platform’s use, and the same open hardware and gel design are equally applicable to validating other acquisition sequences, other imaging modalities, or alternative analysis strategies beyond the fMRI pipeline demonstrated here.

We provide, under open licences in the University of Malta Data Repository (drUM), the full dataset in Brain Imaging Data Structure (BIDS) format; the preprocessing and analysis scripts; a containerised, release-level reproducibility workflow for recomputing and verifying key derivatives; the 3D-printable STL model files for the anatomical phantom; and the tissue-mimicking gel recipes [25]. The reproducibility container is distributed via Docker Hub and archived on Zenodo [26]. Our aim is not only to document this particular phantom but to enable other groups to reproduce it exactly, adapt it to their own scanners and research questions, and improve upon its design, thus lowering the barrier to controlled, dynamic ground-truth experiments and supporting more rigorous, reproducible quality assurance practices across the neuroimaging community.

## Methods

### Anthropomorphic Phantom Construction

#### Anatomical model generation

An anthropomorphic brain-slice model was generated from the *T*_1_-weighted MPRAGE image of a single healthy adult volunteer, acquired on the same scanner used for subsequent phantom imaging (Siemens MAGNETOM Vida 3 T; syngo MR XA20; 64-channel head/neck coil). The anatomical image was processed through the anatomical workflow of fMRIPrep (version 23.0.2[27]), which provided cortical surface reconstructions. White-matter (WM) and grey-matter (GM) surfaces were isolated and converted to multi-material mesh representations using a custom slab-decomposition pipeline. The brain volume was partitioned into six 25 mm axial slabs (in AC–PC-aligned space at 0.5 mm isotropic resolution), and for each slab the following tissue compartments were extracted as separate STL meshes: outer cerebrospinal fluid (CSF), grey matter, white matter, ventricles, and subcortical grey matter. Tissue labels were derived from FreeSurfer parcellations with a partial-volume threshold of 0.5.

A single axial slab (slab 3) was selected for physical fabrication. The model was printed on a Bambu Lab P1S 3D printer using 1.75 *±* 0.03 mm polylactic acid (PLA) filament for the structural elements and polyvinyl alcohol (PVA) filament for dissolvable supports (Bambu Lab, Shenzhen, China). After printing, the model was submerged in warm water with continuous agitation for up to four days until all PVA support material had dissolved, leaving behind the hollow PLA compartment structure (Figure 1).

**Figure 1:**
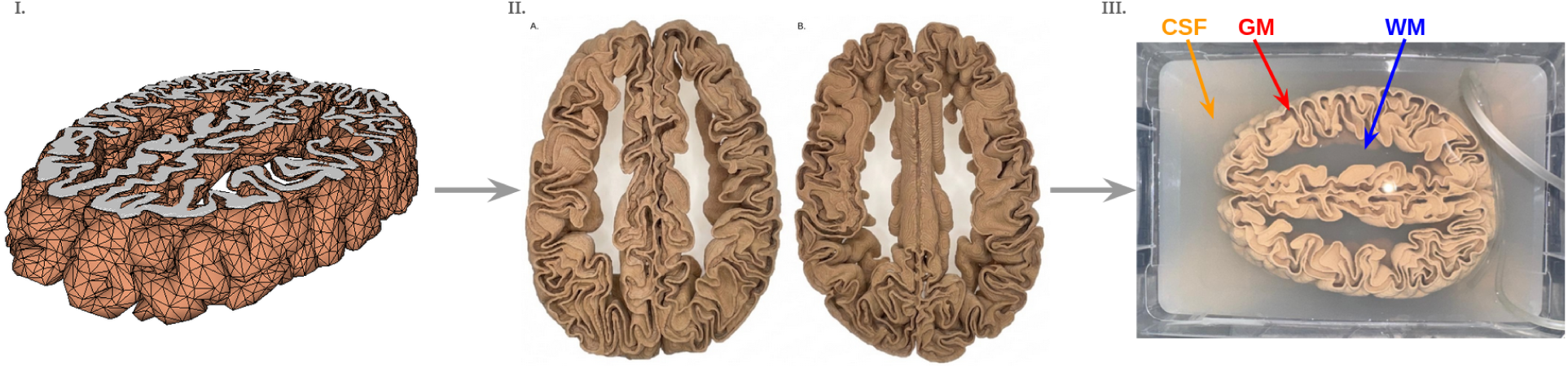
Construction and assembly of the anthropomorphic brain-slice phantom. I. Three-dimensional mesh rendering of axial slab 3. II. The 3D-printed slab after dissolution of the PVA support material, shown from the (A) superior and (B) inferior surfaces. III. Final assembled phantom. The printed slab compartments (WM labelled in blue; GM labelled in red, CSF labelled in orange) were filled with agar-based gels formulated to generate relative MRI contrast resembling that observed between grey matter and white matter. The slab was then embedded in an agar-based gel representing the cerebrospinal fluid compartment. An infusion line was secured to the outer posterior surface of the printed model. During the fMRI runs, hemin and saline solutions were delivered sequentially through the line using a 3 T Tennessee MRI contrast-media injector (Ulrich Medical, Ulm, Germany). Both solutions remained confined to the lumen of the infusion line and did not diffuse into the surrounding gels.

#### Tissue-mimicking gel preparation

Agar-based gels doped with paramagnetic salts were prepared to generate relative MRI contrast resembling that observed between human WM, GM, and CSF. The base components were European bacteriological agar (Condalab, Madrid, Spain), sodium chloride (NaCl, 99.5%; ThermoScientific, Denmark), nickel (II) chloride hexahydrate (NiCl_2_, 98%; ThermoScientific, France), and manganese (II) chloride anhydrous (MnCl_2_, *>*98%; Fluka Chemika, Buchs, Switzerland). NiCl_2_ and MnCl_2_ were selected as paramagnetic relaxation modifiers following established precedent in the MRI phantom literature, where nickel- and manganese-doped agarose gels have been widely used to generate independently tunable *T*_1_ and *T*_2_ contrast [28–30]. Gel compositions are summarised in Table 1.

**Table 1:** Tissue-mimicking gel compositions.

| Tissue | Composition |
| --- | --- |
| White matter | 1% (w/v) agar + 0.66% (w/v) NaCl + 1.7 mM $\text{NiCl}_2$ + 0.085 mM $\text{MnCl}_2$ |
| Grey matter | 1.5% (w/v) agar + 0.4 mM $\text{NiCl}_2$ |
| Cerebrospinal fluid | 0.5% (w/v) agar |

Each gel was prepared by suspending the components in the appropriate proportions in lukewarm deionised water, then covering with aluminium foil to reduce evaporation, and heating on a hot plate with constant stirring until the agar had fully dissolved. The solutions were maintained at a sustained boil for two minutes, transferred to a 60 *^◦^*C incubator for 30 minutes (to facilitate degassing and thermal equilibration), and finally allowed to cool at room temperature to approximately 50 *^◦^*C before use.

#### Phantom assembly

The SiGn anthropomorphic phantom was assembled in a 3 L rectangular plastic container. A 1.5 L base layer of CSF-composition gel was poured and allowed to solidify fully. The 3D-printed anatomical model was positioned on the base layer and secured with a thin secondary layer of CSF gel; the assembly was left to solidify overnight at 4 *^◦^*C. WM and GM gels were then sequentially injected into the corresponding model compartments using separate syringes. The infusion line was positioned on the outer posterior surface of the model, external to the printed cortical geometry. This was done primarily for practical reasons: this location simplified fabrication and line routing, and avoided introducing a foreign object into the gel-filled tissue compartments where it could disrupt the intended WM/GM contrast.

Confining hemin delivery to the lumen of the infusion line, rather than injecting it directly into the white or grey matter gel, was also intended to preserve a reproducible, well-characterised source geometry: the solution’s spatial extent and timing remain fully controlled by the infusion system rather than being subject to the unpredictable diffusion and absorption properties of the surrounding gel. We note that, unlike a human head, the phantom has no skull or scalp to remove computationally; there is consequently no brain-extraction step analogous to human preprocessing that could remove this signal from the analysis. The present study’s registration and normalisation pipeline instead treats the phantom’s full geometry, including the CSF-mimicking gel in which the infusion line is embedded, as the object of interest, so the posterior activation remains fully within the analysed field of view. After overnight solidification at 4 *^◦^*C, the entire model was submerged in a final layer of CSF-composition gel to a depth of approximately 2 cm above the top surface of the model. The phantom was left to solidify at room temperature before scanning (Figure 1).

#### Signal-altering contrast agent

Hemin (C_34_H_32_ClFeN_4_O_4_, Sigma-Aldrich, St. Louis, MO, USA), the chloride salt of oxidised haem, was used as a paramagnetic contrast agent. Dissolution in alkaline solution converts hemin to hematin, a paramagnetic substance that shortens 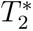 (and, to a lesser extent, *T*_1_ and *T*_2_) relaxation times, thereby mimicking the susceptibility effect of deoxyhaemoglobin exploited in BOLD fMRI. Hemin stock solutions were prepared at concentrations of 10 mg/mL, 0.78 mg/mL, and 0.39 mg/mL by dissolving hemin in deionised water with dropwise addition of 1 M sodium hydroxide (NaOH, Sigma-Aldrich, St. Louis, MO, USA) until full solubility was achieved [31].

### MRI Data Acquisition

All imaging was performed on a Siemens MAGNETOM Vida 3 T scanner (syngo MR XA20) using a 64-channel head/neck receive coil (HeadNeck 64 CS). Data were acquired across two scanning sessions.

#### Structural imaging

A *T*_1_-weighted anatomical image of the phantom was acquired in each session using a 3D MPRAGE sequence (TR = 2190 ms; TE = 2.66 ms; TI = 925 ms; flip angle = 8*^◦^*; 1.0 mm isotropic voxels; matrix = 256 *×* 176 *×* 256).

#### Infusion paradigm design

A 3 T Tennessee MRI contrast-media injector (Ulrich Medical, Ulm, Germany) was used to deliver saline (Sodium Chloride 0.9% Solution for Infusion [Viaflo], Baxter, Sintra, Portugal) and freshly prepared hemin solutions through the lumen of an infusion tube adhered to the outer posterior surface of the PLA model. The solutions remained confined to the tube lumen and did not enter or diffuse into the surrounding gels. The injector was equipped with two fluid inputs: a saline bag and a syringe.

The infusion schedules consisted of alternating periods of saline and hemin delivery. For the long-block paradigms, the solutions were delivered at a flow rate of 0.5 mL/s. The Session 1 long-block paradigm comprised four 22-s hemin-delivery periods, corresponding to 11 mL per hemin period, separated by 42-s saline-delivery periods, corresponding to 21 mL per saline period. The Session 2 long-block paradigm used the same block durations and flow rate.

For the Session 1 short-block paradigm, the flow rate was increased to 1 mL/s. This paradigm comprised two 21-s hemin-delivery periods, corresponding to 21 mL per hemin period, separated by 42-s saline-delivery periods, corresponding to 42 mL per saline period. These schedules were selected to produce repeated, temporally distinct signal changes within the available acquisition duration while accommodating fluid transit and washout. The long- and short-block arrangements were included to assess whether the phantom-generated signal could be detected under different temporal delivery conditions relevant to block-design fMRI. The programmed infusion schedules and corresponding convolved GLM regressors are shown in Figure 2.

**Figure 2:**
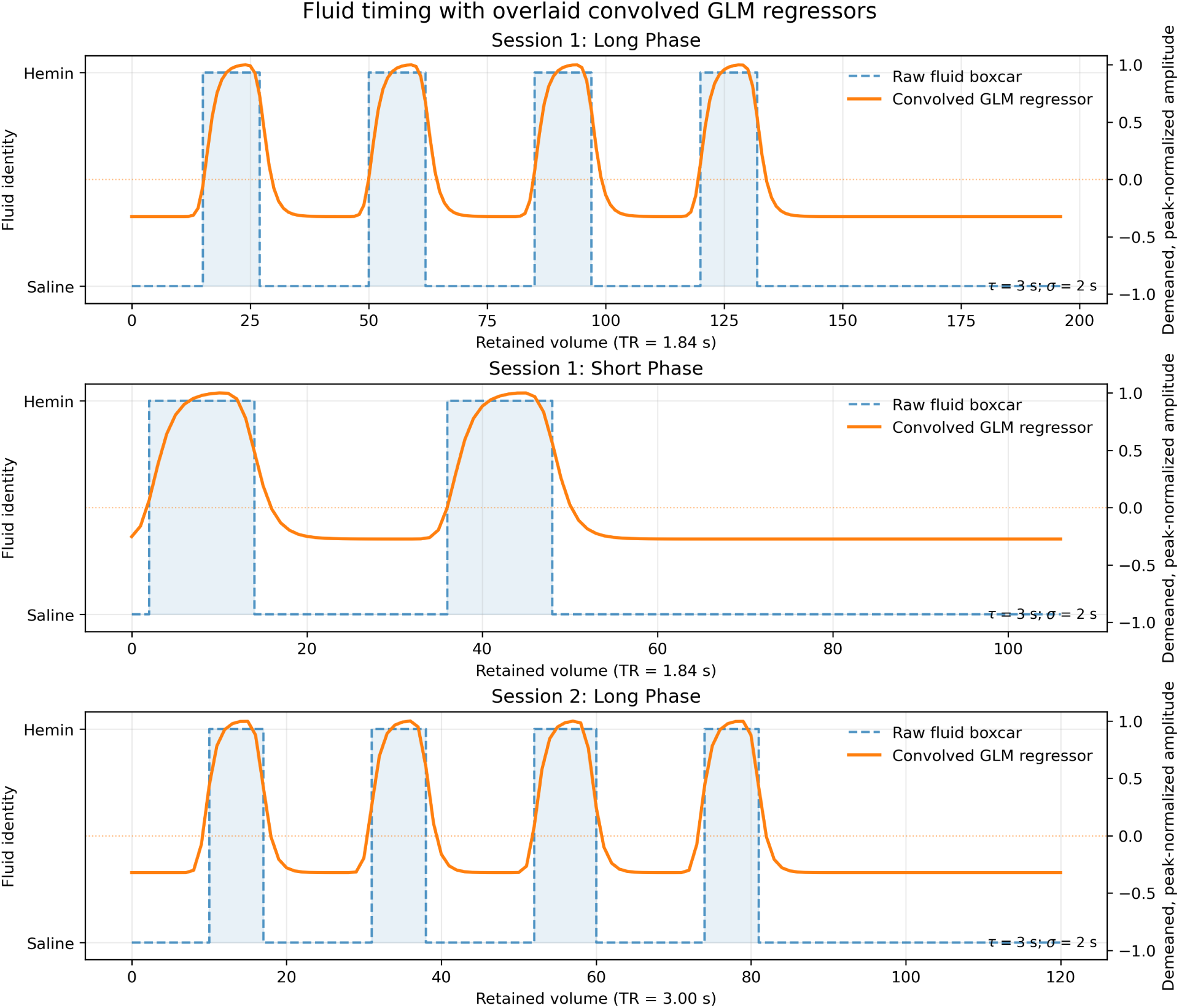
Infusion timing paradigms and corresponding convolved GLM regressors. The upper and middle panels show the Session 1 long-block paradigm (200 volumes; TR = 1.84 s), containing four hemin-delivery periods, and the short-block paradigm (110 volumes; TR = 1.84 s), containing two hemin-delivery periods, respectively. The lower panel shows the Session 2 long-block paradigm (123 volumes; TR = 3.00 s), containing four hemin-delivery periods. In each panel, the dashed boxcar and shaded regions indicate the programmed switches between saline and hemin delivery, while the solid line shows the corresponding fluid regressor used in the general linear model. The regressor was generated by convolving the boxcar with a causal exponential washout kernel (*τ* = 3 s), followed by Gaussian temporal smoothing (*σ* = 2 s), downsampling to the acquisition repetition time, demeaning, and peak normalization. The saline-only control followed the Session 1 long-block timing paradigm, but both injector inputs contained saline.

#### Functional imaging session 1

Five functional runs were acquired using a 2D gradient-echo EPI sequence (TR = 1840 ms; TE = 20 ms; flip angle = 65*^◦^*; 2.5 mm isotropic voxels; matrix = 96 *×* 96; 60 slices; multiband acceleration factor = 4; phase-encoding direction H*→*F; total readout time = 55.1 ms; effective echo spacing = 0.58 ms). Four runs evaluated two hemin concentrations (0.78 and 0.39 mg/mL), with each concentration tested using both the long- and short-block paradigms described above. The long- and short-block runs comprised 200 and 110 volumes, respectively. During these runs, the active injector input alternated between the saline bag and the hemin-containing syringe according to the programmed infusion schedules shown in Figure 2.

A fifth run followed the long-block paradigm and served as a saline-only control. In this run, both injector inputs contained saline, allowing signal fluctuations associated with switching between injector inputs to be assessed independently of hemin delivery. A reverse-phase-encoded gradient-echo EPI reference acquisition (10 volumes, F*→*H) was acquired for distortion correction; `topup` was estimated from its first volume together with the first volume of the first BOLD run.

#### Functional imaging session 2

A single task condition was acquired at a hemin concentration of 10 mg/mL using a long-block paradigm (123 volumes) as described in Figure 2. The EPI sequence parameters differed from Session 1 to reduce the T1 and flow effects noted in Session 1 (TR = 3000 ms; TE = 20 ms; flip angle = 40*^◦^*; 2.5 mm isotropic voxels; 96 *×* 96 matrix; 60 slices; phase-encoding direction F*→*H; total readout time = 26.6 ms; effective echo spacing = 0.28 ms). Two distortion-corrected outputs were produced from this run using FSL `topup`: one estimated from a reverse-phase-encoded gradient-echo EPI pair and one estimated from a reverse-phase-encoded spin-echo EPI pair (SE EPI: TR = 4700 ms; TE = 48 ms; flip angle = 90*^◦^*), with opposing phase-encode directions (F*→*H and H*→*F).

### Preprocessing

Preprocessing was performed primarily using FSL 6.0.7[32], with ancillary tooling for data conversion and optional spatial normalisation steps. DICOMs were converted using dcm2niix (v1.0.20250505)[33] and data were organised into the BIDS format[34]. The human T1w scan was converted using dcm2niix (v1.0.20190902) due to it having been preprocessed for a previous project.

#### Structural processing and registration

Three coordinate systems are used throughout the released derivatives, each identified by a BIDS `space-` entity. `space-ACPC` denotes the human participant’s AC–PC-aligned *T*_1_w grid; this is the reference for all human anatomical files (`sub-human`) and for any phantom file that has been resampled into the human’s coordinate frame. `space-PhantomT1w` denotes the reoriented native grid of the phantom’s own session-specific *T*_1_w image; BOLD reference images coregistered to the phantom anatomy carry this label. `space-MNI152NLin2009cAsym` denotes the standard MNI template space, reached by composing the `PhantomT1w` *→* `ACPC` rigid transform with the participant’s previously computed `ACPC` *→* MNI affine and nonlinear warps [35, 36]. The same three labels are used consistently in filenames and in the `from-`/`to-` entities of the corresponding transform files.

#### Susceptibility distortion correction

EPI susceptibility-induced distortions were estimated and corrected using FSL’s `topup`[37]. In Session 1, `topup` was estimated from the reverse-phase-encoded gradient-echo EPI pair (F*→*H and H*→*F; readout time = 55.1 ms) and the resulting field was applied to all five functional runs via `applytopup` (Jacobian modulation). Each corrected BOLD time-series was then reoriented to standard orientation using the same axis swap applied to the BOLD data. In Session 2, two separate `topup` estimates were computed: one from gradient-echo (Gre) EPI pairs and one from spin-echo (Se) EPI pairs (readout time = 26.6 ms), yielding two independently corrected versions of the functional data. Both corrected outputs were reoriented as above.

#### Functional-to-structural coregistration

For each distortion-corrected BOLD run, a temporal mean image was computed and used as the reference volume (boldref) for rigid-body coregistration (6 DOF, normalised mutual information; FSL FLIRT) to the session’s reoriented phantom *T*_1_w (`space-PhantomT1w`). A chained transformation from BOLD native space to the human participant’s AC–PC-aligned *T*_1_w (`space-ACPC`) was then computed by concatenating the boldref *→* `PhantomT1w` and `PhantomT1w` *→* `ACPC` transforms using `convert xfm`.

### Statistical Analysis

Voxelwise activation mapping was performed using a general linear model (GLM) implemented through FSL’s FILM with local autocorrelation correction[38]. Analyses were conducted independently for each functional run.

#### Temporal preprocessing

To ensure steady-state longitudinal magnetisation, the first 5 s of each functional run were discarded prior to analysis (corresponding to 3 volumes for TR = 1.84 s and 2 volumes for TR = 3.0 s, rounded up to the nearest whole TR). A high-pass temporal filter with a cutoff period of 100 s was applied. This cutoff was selected to attenuate low-frequency scanner drift while preserving signal at the frequencies represented in the infusion paradigm, consistent with commonly used FSL-based fMRI analysis settings [38].

#### Spatial preprocessing

Functional data were spatially smoothed with a Gaussian kernel of 6 mm FWHM (*σ ≈* 2.55 mm), a value representative of typical smoothing kernels used in standard fMRI preprocessing pipelines, sufficient to improve signal-to-noise ratio while remaining smaller than the phantom’s compartment dimensions.

#### GLM design

Event timing was derived from BIDS `events.tsv` files, with onset times adjusted by the trimmed offset so that regressors aligned with the retained data. The design matrix included three regressors: (1) a *fluid* regressor modelling the identity of the fluid present at the phantom (hemin or saline), generated by convolving the infusion timing with a washout kernel (exponential decay, *τ* = 3 s) and smoothing with a Gaussian (*σ* = 2 s) to approximate the temporal profile of contrast-agent arrival and clearance at the imaging voxel. The washout kernel time constant (*τ* = 3 s) was selected to approximate the expected transit and clearance time of contrast agent through the infusion line and surrounding gel at the flow rates used, based on preliminary timing observations during phantom testing; (2) a *switch* regressor modelling the brief transient at each pump changeover, when the infusion line switched from one substance to the other; and (3) a *white matter* confound regressor extracted from a WM mask derived from the 3D-printed compartment geometry, propagated from the source human anatomical space into functional space. The fluid and switch regressors were generated at a high temporal resolution of 0.1 s, then downsampled to the acquisition TR; the WM confound was extracted directly from the trimmed functional data. All non-intercept regressors were demeaned and peak-normalised to unit magnitude before model fitting. The primary contrast of interest tested the *fluid* regressor against baseline.

Statistical maps were thresholded at *Z >* 3.1, and cluster-level significance was assessed at *p <* 0.05 (corrected) using Gaussian random field theory, with smoothness estimated from the residual images via FSL’s smoothest.

#### Sensitivity analyses

To assess the dependence of the GLM results on selected analysis parameters, the models were re-estimated while varying one parameter at a time and holding all remaining settings at their reference values. Spatial-smoothing sensitivity was evaluated using Gaussian kernels of 0, 3, 6, and 8 mm FWHM, with the high-pass cutoff fixed at 100 s and the exponential washout time constant fixed at *τ* = 3 s. Temporal-filter sensitivity was evaluated using high-pass cutoffs of 50, 100, 150, and 200 s, with spatial smoothing fixed at 6 mm FWHM and *τ* = 3 s. Sensitivity to the assumed washout model was evaluated using exponential time constants of *τ* = 1, 3, 6, and 10 s, with spatial smoothing fixed at 6 mm FWHM and the high-pass cutoff fixed at 100 s. The temporal Gaussian smoothing of the fluid regressor was retained at *σ* = 2 s throughout.

The released GLM workflow reads the spatial-smoothing kernel, temporal high-pass cutoff, exponential washout time constant, and temporal Gaussian smoothing parameter from the central `config.env` file. Alternative values can therefore be evaluated by modifying the corresponding configuration entry and rerunning `code/scripts/03_run_glm_v1_0_0.sh` on the supplied preprocessed data.

#### Spatial normalisation of statistical maps

Unthresholded and thresholded *Z*-statistic maps were warped from BOLD native space into space-PhantomT1w (the phantom’s own native *T*_1_w grid) and, where the corresponding transforms were available, into space-ACPC (the human participant’s AC–PC-aligned *T*_1_w) and space-MNI152NLin2009cAsym (the MNI template), using the precomputed rigid-body and nonlinear transformations described above.

#### Independent component analysis

Probabilistic independent component analysis (ICA) was also performed on each run using FSL’s MELODIC, with automatic dimensionality estimation (background threshold = 3; mixture-model threshold = 0.5).

### Software and Data Availability

Preprocessing and analysis pipelines were implemented as versioned shell scripts, with SHA-256 checksums recorded for key scripts, configuration files, and selected inputs and outputs as part of the provenance and verification workflow. The analysis code used Python 3.10.12 with NumPy 1.26.4, SciPy 1.15.3, NiBabel 5.4.2, and Matplotlib 3.10.8 alongside FSL 6.0.7. Raw and derivative data were organised following the Brain Imaging Data Structure (BIDS) specification. The full dataset, processing scripts, and reproducibility workflow are available via drUM at https://doi.org/10.60809/drum.31411158. The corresponding container image is distributed via Docker Hub and archived on Zenodo at https://doi.org/10.5281/zenodo.19495290. The image additionally bundles third-party software governed by its own upstream licenses, which remain in force for those components.

### Use of Large Language Models

Data organisation, scripting, and writing polishing were aided using multiple large language models (LLMs), including Claude, ChatGPT, and Gemini. All AI-assisted outputs were reviewed, verified, and edited by the authors before inclusion in the final workflow and manuscript.

## Results

The complete preprocessing and analysis pipeline, from BIDS-formatted raw data through susceptibility distortion correction, coregistration, spatial normalisation, and GLM-based statistical mapping, was successfully executed for all functional runs across both scanning sessions. The pipeline, together with the BIDS-formatted dataset, 3D-printable STL files, gel recipes, and containerised reproducibility workflow, has been made openly available at https://doi.org/10.60809/drum.31411158. This release enables other groups to replicate the full workflow on their own hardware or to verify the derivatives reported here.

The anthropomorphic geometry of the SiGn phantom enabled the successful integration of phantom data into standard fMRI preprocessing pipelines. The *T*_1_-weighted image of the phantom was successfully registered into the human participant’s AC–PC space (`space-ACPC`; Figure 3) and into MNI standard space (`space-MNI152NLin2009cAsym`; Figure 4).

**Figure 3:**
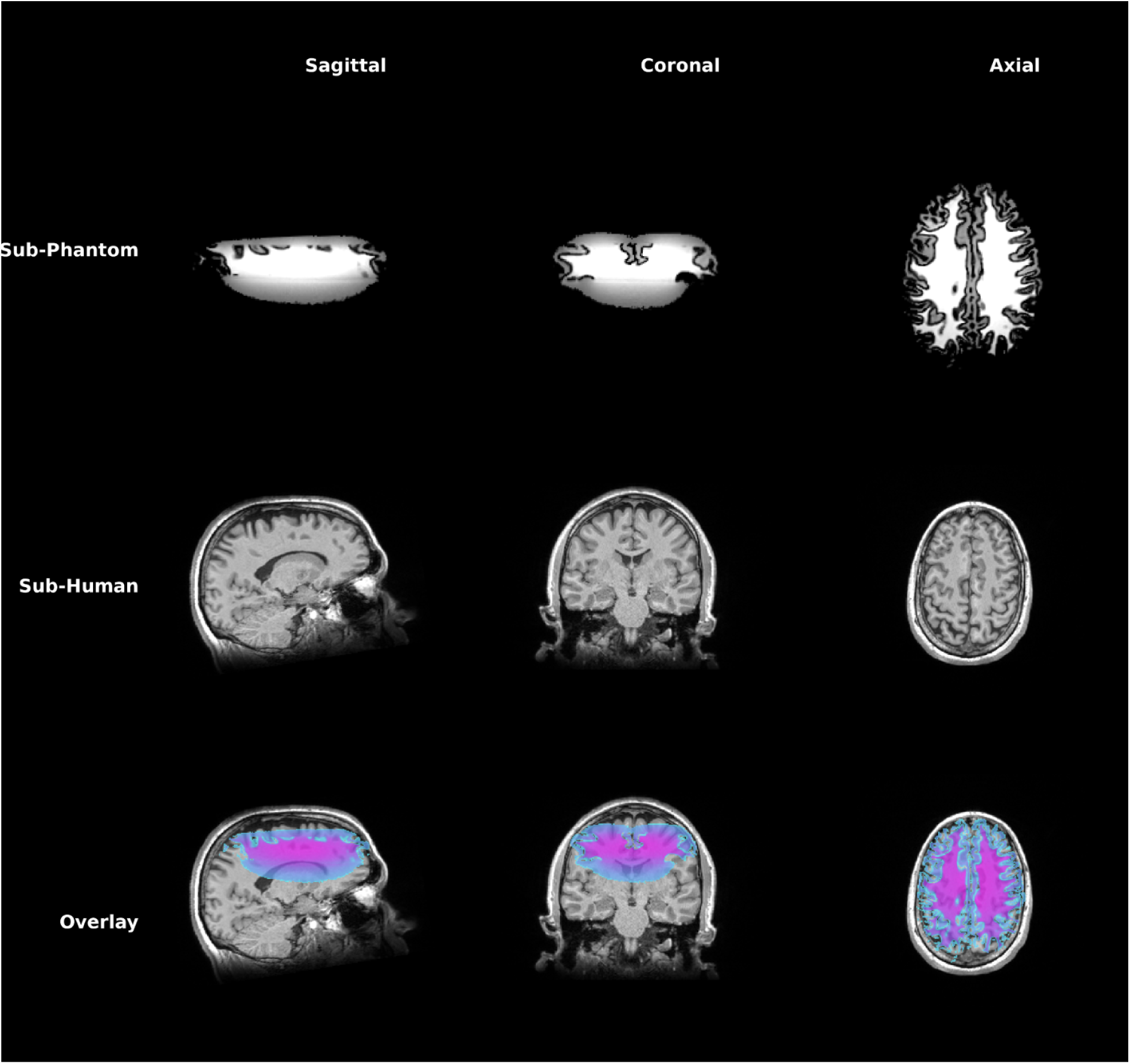
Spatial registration of the anthropomorphic brain phantom to the human anatomical image. Representative sagittal, coronal, and axial views are shown. The upper row shows the Session 1 *T*_1_-weighted MR image of the SiGn phantom. The middle row shows the *T*_1_-weighted MR image of the human reference anatomy. The lower row shows the spatially registered phantom MR image, displayed using the cool colormap, overlaid on the human anatomical MR image.

**Figure 4:**
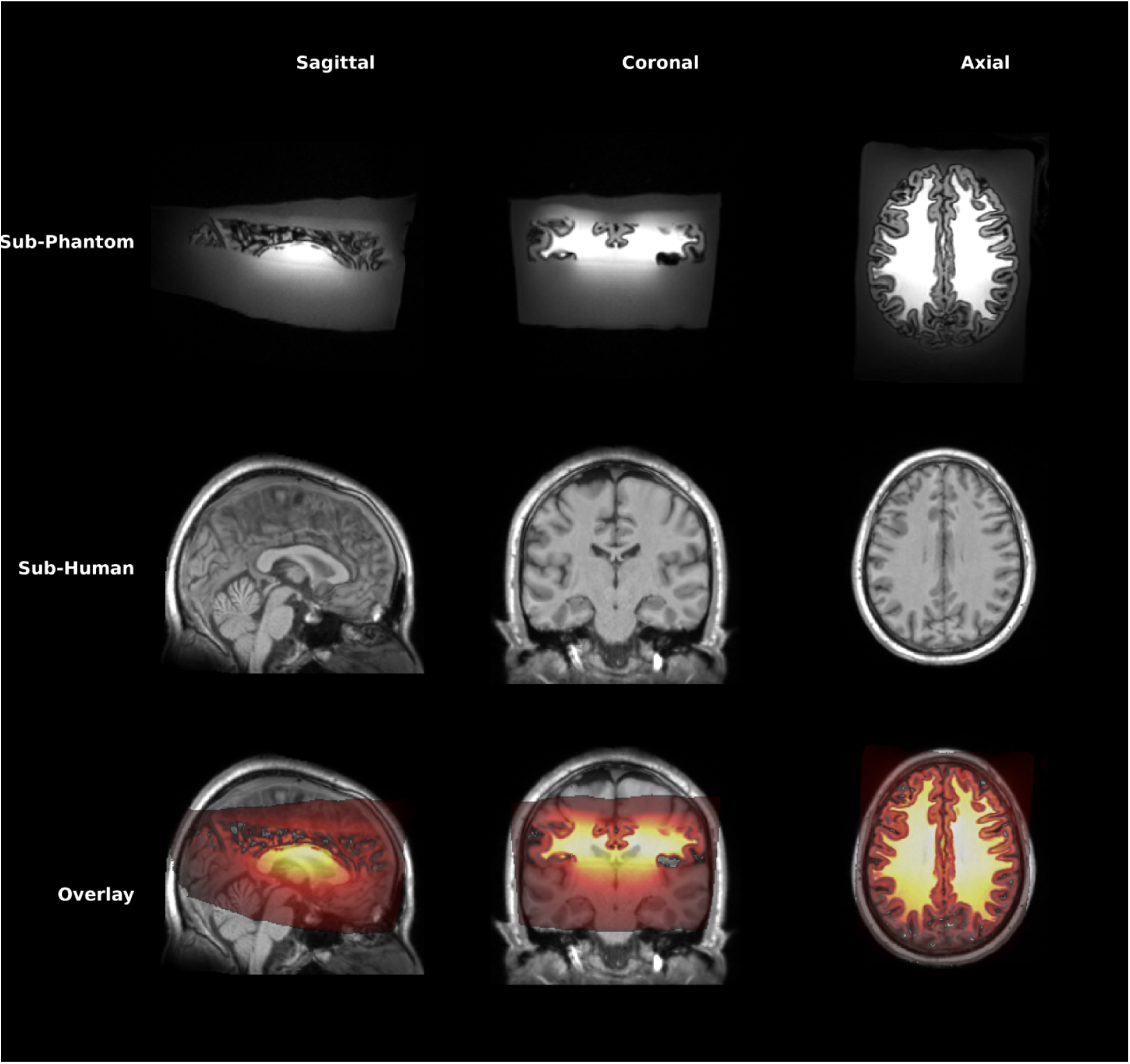
Spatial registration of the anthropomorphic brain phantom and human reference anatomy to MNI152 standard space. Representative sagittal, coronal, and axial views are shown. The upper row shows the Session 1 *T*_1_-weighted MR image of the SiGn phantom after registration to MNI152 space. The middle row shows the Session 1 *T*_1_-weighted human reference MR image in MNI152 space. The lower row shows the registered phantom MR image, displayed using the hot colormap, overlaid on the registered human anatomical image to illustrate their spatial correspondence.

To validate the phantom’s capacity to generate detectable, spatially localised signal changes, voxelwise GLM analyses were performed independently for each functional run. The primary contrast tested the *fluid* regressor, which modelled the identity of the solution present at the phantom, against baseline.

Figure 5 illustrates the GLM results for the hemin-saline runs of both Sessions 1 and 2, and the saline-control run of Session 1. In Session 1, hemin infusion at a concentration of 0.78 mg/mL produced significant activation clusters in both the long-block (Figure 5, first row, left) and short-block (Figure 5, second row, left) paradigms. The 0.39 mg/mL hemin concentration also produced significant clusters in both the long-block (Figure 5, first row, right) and short-block (Figure 5, second row, right) paradigms, with comparatively reduced magnitude and spatial extent in the short-block condition. In the hemin conditions, dominant clusters were spatially concentrated in the posterior region of the phantom, consistent with the known position of the infusion tube.

**Figure 5:**
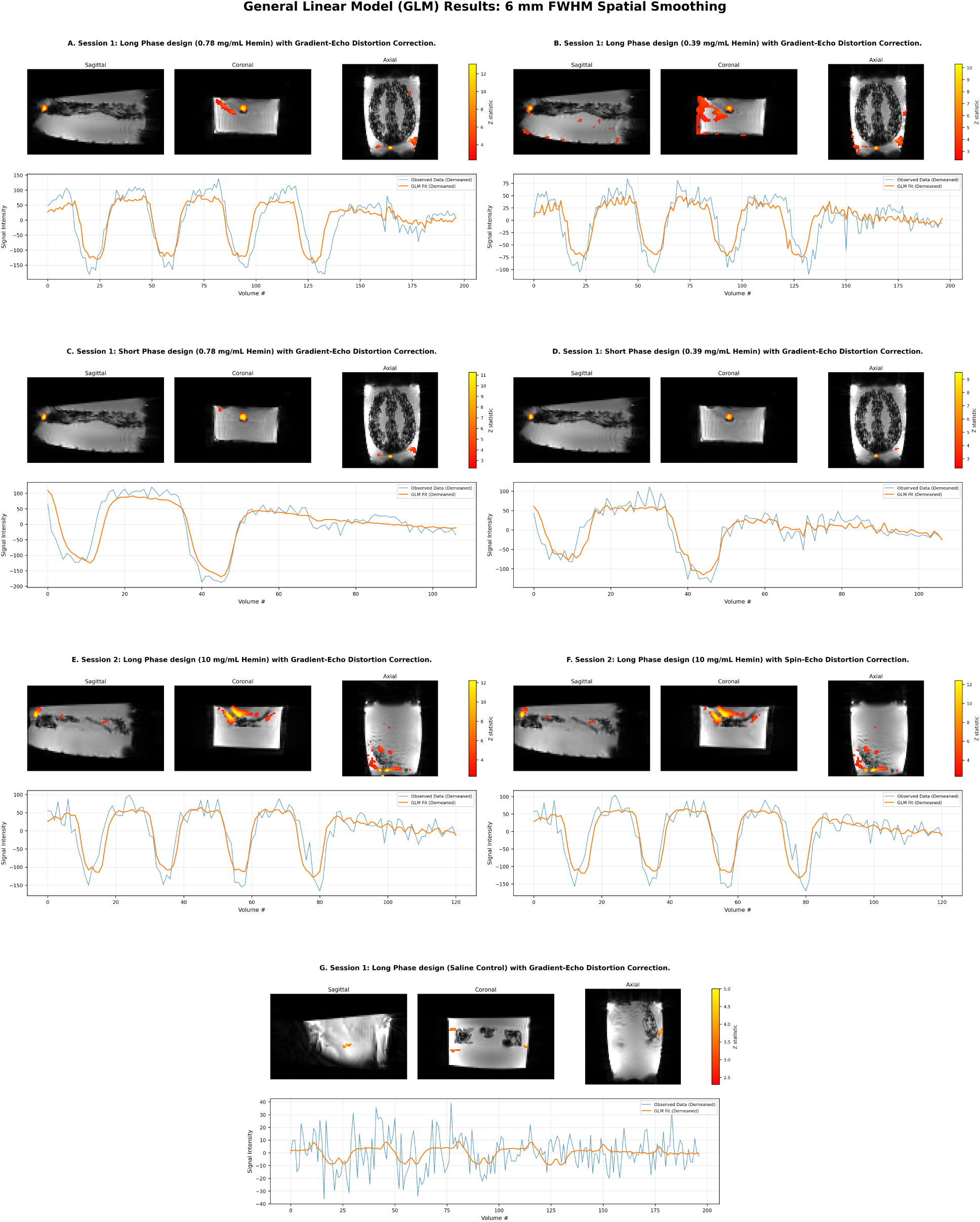
General Linear Model (GLM) results for the hemin-saline and saline-control runs. Each panel shows thresholded GLM statistical maps in representative sagittal, coronal, and axial views (upper portion), together with the demeaned signal-intensity time course from the analysed region and the corresponding demeaned GLM fit (lower portion). The first row shows the Session 1 long-phase runs acquired with 0.78 mg/mL hemin (left) and 0.39 mg/mL hemin (right) after gradient-echo (Gre) distortion correction. The second row shows the corresponding Session 1 short-phase runs acquired with 0.78 mg/mL hemin (left) and 0.39 mg/mL hemin (right) after Gre distortion correction. The third row shows the Session 2 long-phase run acquired with 10 mg/mL hemin after distortion correction based on Gre EPI images (left) and spin-echo (Se) EPI images (right). The bottom panel shows the Session 1 long-phase saline-only control after Gre distortion correction. In the time-course plots, the observed demeaned signal is shown in blue and the demeaned GLM fit in orange; in the statistical maps, the colour bars indicate the magnitude of the GLM test statistic.

The saline-only control condition showed weaker and less specific effects than the hemin conditions, with a lower peak statistic and false activation outside the expected zone (Figure 5, fourth row). Relative to saline, the hemin runs produced substantially stronger and more spatially focused activation in the expected posterior region.

In Session 2, which employed a hemin concentration of 10 mg/mL with modified acquisition parameters (TR = 3000 ms; flip angle = 40*^◦^*), significant activation was again detected in the posterior region using both gradient-echo-based (Figure 5, third row, left) and spin-echo-based (Figure 5, third row, right) susceptibility distortion correction. The time-series data at the peak voxels showed clear modulation time-locked to the infusion paradigm in these conditions. Inspection of the time-series data and independent component analyses indicated that signal changes were not uniform across the infusion region (Figure 5, coronal views). Voxels within the core of the infusion tube showed a signal change of opposite polarity to the surrounding gel, with an increase rather than the expected decrease in signal intensity during hemin delivery. This inverted response is consistent with *T*_1_ and inflow effects: fresh, fully magnetised solution entering the imaging volume produces a transient signal enhancement that opposes the 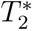-mediated darkening in adjacent tissue-mimicking gel. Although the modified acquisition parameters in Session 2 were designed to attenuate these contributions, this effect persisted, but at reduced magnitude.

The signal mechanism underlying the phantom differs fundamentally from the neurovascular BOLD effect. The observed 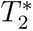 modulation arises from the flow and diffusion of a paramagnetic contrast agent through a tube embedded in agar gel, rather than from metabolically coupled changes in blood oxygenation. The phantom’s signal changes should therefore be interpreted as a controlled, physically grounded surrogate for fMRI activation, sufficient to test whether standard acquisition and analysis pipelines can correctly detect and localise time-locked signal changes within an anatomically realistic geometry, rather than as a faithful model of haemodynamic responses.

## Discussion

This work describes the development, and initial validation of the SiGn anthropomorphic brain phantom, together with a fully open pipeline for preprocessing and analysing the resulting data. The principal contribution is not the phantom itself but the complete, reproducible workflow, from 3D-printable anatomy and tissue contrast-mimicking gel recipes through BIDS-formatted acquisitions to containerised analysis code, that enables other groups to fabricate, scan, and analyse an equivalent device on their own hardware.

The GLM analyses confirmed that hemin infusion produced detectable, spatially localised signal changes in the outer posterior region of the phantom, consistent with the known position of the infusion line. These changes were concentration-dependent and substantially stronger than those observed in the saline control condition, supporting the interpretation that the dominant effects reflect genuine 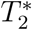-shortening by the paramagnetic contrast agent rather than scanner drift or other systematic artefacts. However, inspection of the raw time-series data and the independent component analyses revealed a signal change of opposite polarity within the core of the infusion tube. This inverted response is most parsimoniously explained by *T*_1_ effects and inflow-related signal enhancement: fresh solution entering the tube carries full longitudinal magnetisation that has not experienced prior radiofrequency excitation, producing a transient signal increase that opposes the 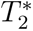-mediated darkening in the surrounding gel. The longer repetition time and reduced flip angle adopted in Session 2 were intended to attenuate these contributions, and the persistence of the effect suggests that a complete separation of *T*_1_ and 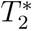 components will require either dedicated multi-echo acquisitions or flow-compensation strategies in future iterations of the phantom.

Despite these confounds in the tube lumen, the critical observation is that the 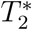-weighted signal changes in the gel surrounding the infusion site behaved as expected: they were temporally locked to the infusion paradigm, spatially confined to the region of contrast-agent diffusion, and amenable to standard GLM-based detection. This establishes the SiGn phantom as a viable platform for generating controlled, ground-truth activation signals within an anatomically realistic geometry.

The dependence of the GLM results on the selected analysis parameters was assessed by re-estimating the models while varying one parameter at a time and holding all other settings at their reference values. For each parameter setting, all five Session 1 runs and both distortion-corrected versions of the Session 2 run were analysed using the same design matrix, confound-regression strategy, contrast definition, and statistical-thresholding procedure as the baseline analysis. The resulting statistical maps and fitted time courses were then compared qualitatively across parameter values. Figures 6 to 8 illustrate these comparisons for the Session 1 short-phase condition with 0.39,mg/mL hemin. These analyses were intended to characterise the sensitivity of the results to processing and modelling choices rather than to identify an optimal parameter combination. The predefined settings of 6,mm FWHM spatial smoothing, a 100,s high-pass cutoff, and an exponential washout time constant of 3,s produced interpretable results and were therefore retained for the baseline analysis. Nevertheless, the comparisons demonstrate that parameter selection can influence statistical detectability, spatial extent, and model fit. Consequently, alternative parameter settings may be more appropriate depending on the intended application.

**Figure 6:**
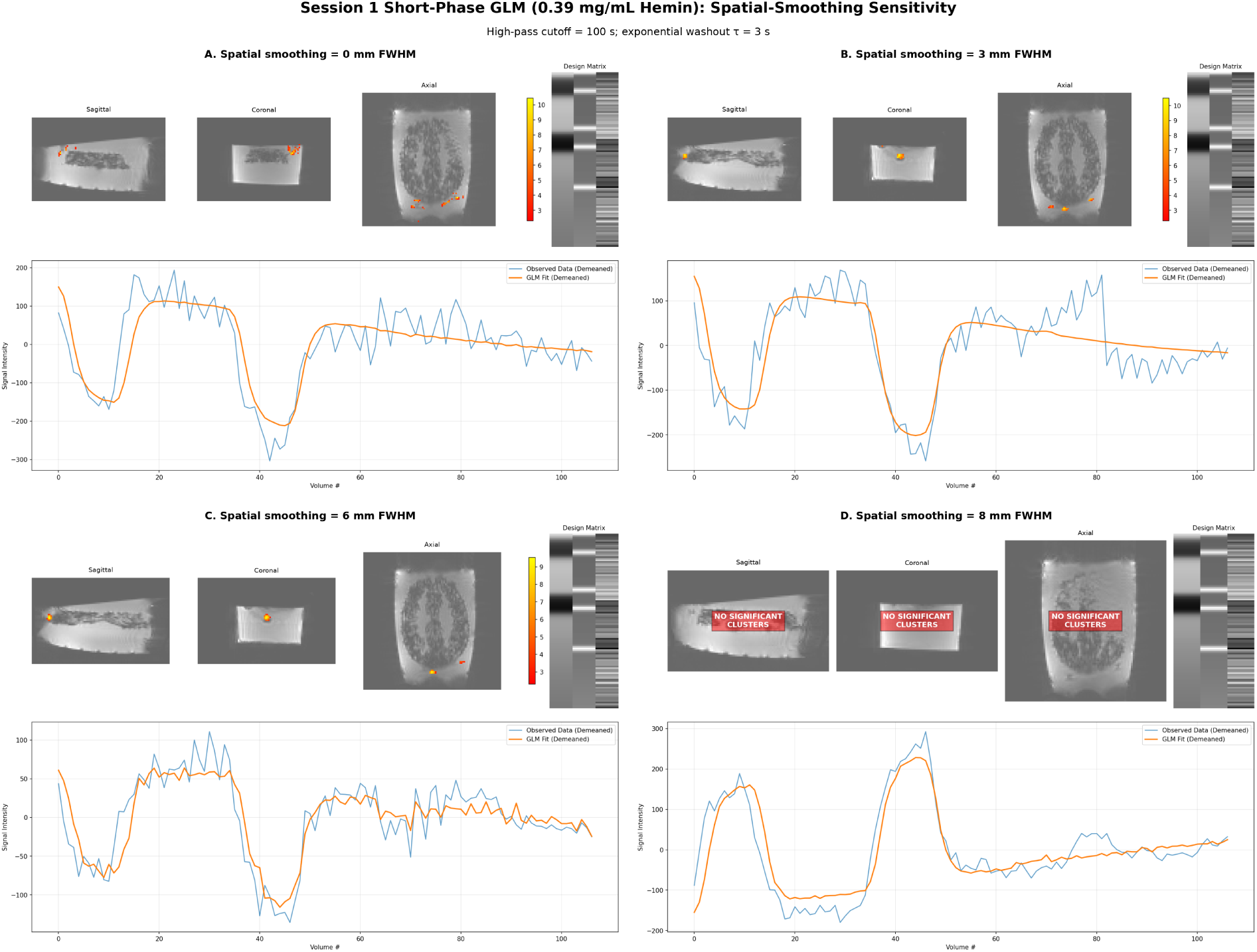
Sensitivity of the Session 1 short-phase 0.39 mg/mL hemin GLM result to spatial smoothing. Panels show results obtained using Gaussian smoothing kernels of (A) 0 mm, (B) 3 mm, (C) 6 mm, and (D) 8 mm FWHM, while the high-pass cutoff and exponential washout time constant were held fixed at 100 s and *τ* = 3 s, respectively. Each panel presents thresholded GLM *Z*-statistic maps in representative sagittal, coronal, and axial views, together with the demeaned observed signal time course (blue) and demeaned GLM fit (orange). Where no cluster survived correction, this is indicated on the anatomical views and the time course was evaluated at the uncorrected statistical peak. All images were derived from the gradient-echo-distortion-corrected functional data.

**Figure 7:**
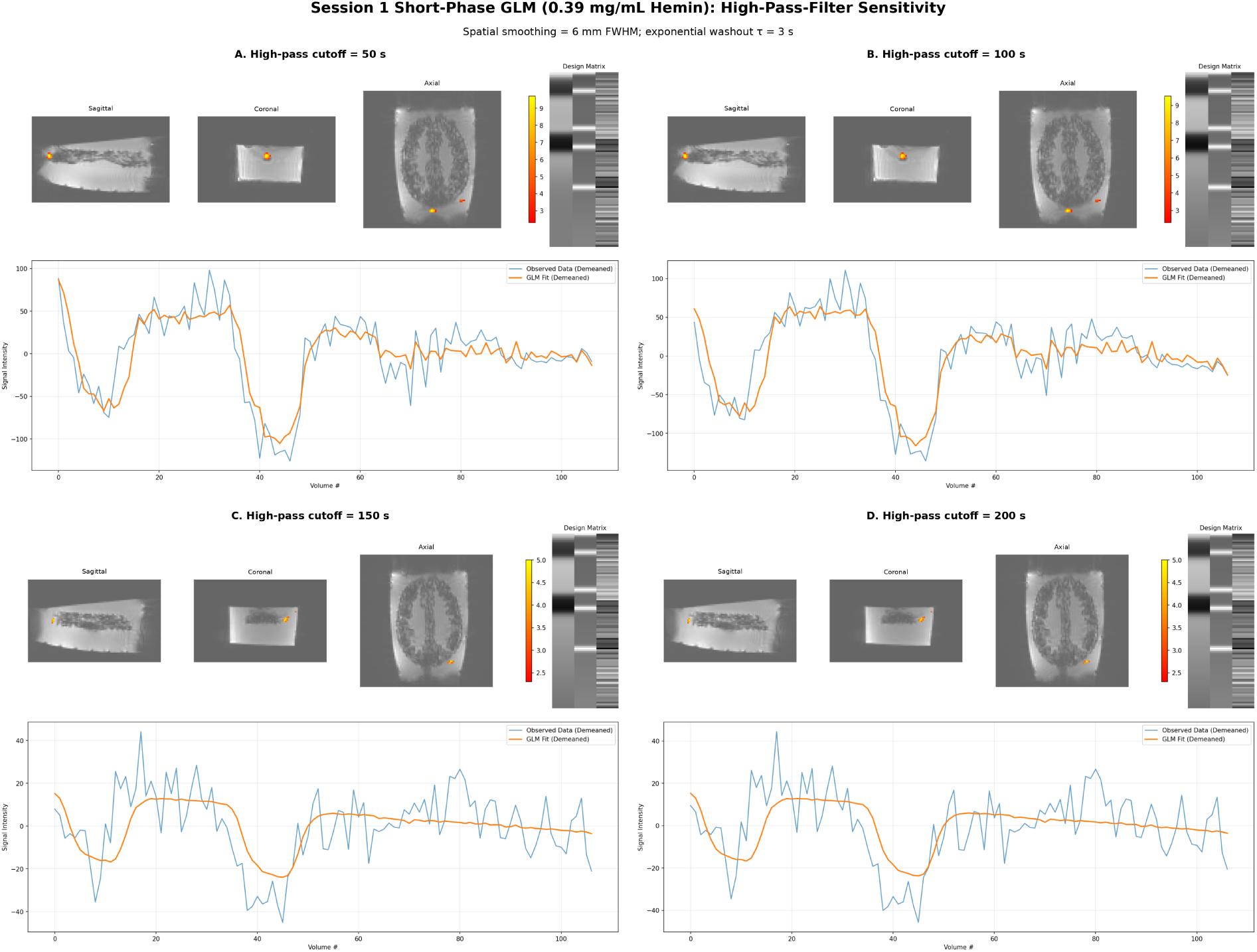
Sensitivity of the Session 1 short-phase 0.39 mg/mL hemin GLM result to temporal high-pass filtering. Panels show results obtained using high-pass cutoff periods of (A) 50 s, (B) 100 s, (C) 150 s, and (D) 200 s, while spatial smoothing and the exponential washout time constant were held fixed at 6 mm FWHM and *τ* = 3 s, respectively. Each panel presents thresholded GLM *Z*-statistic maps in representative sagittal, coronal, and axial views, together with the demeaned observed signal time course (blue) and demeaned GLM fit (orange). Where no cluster survived correction, this is indicated on the anatomical views and the time course was evaluated at the uncorrected statistical peak. All images were derived from the gradient-echo-distortion-corrected functional data.

**Figure 8:**
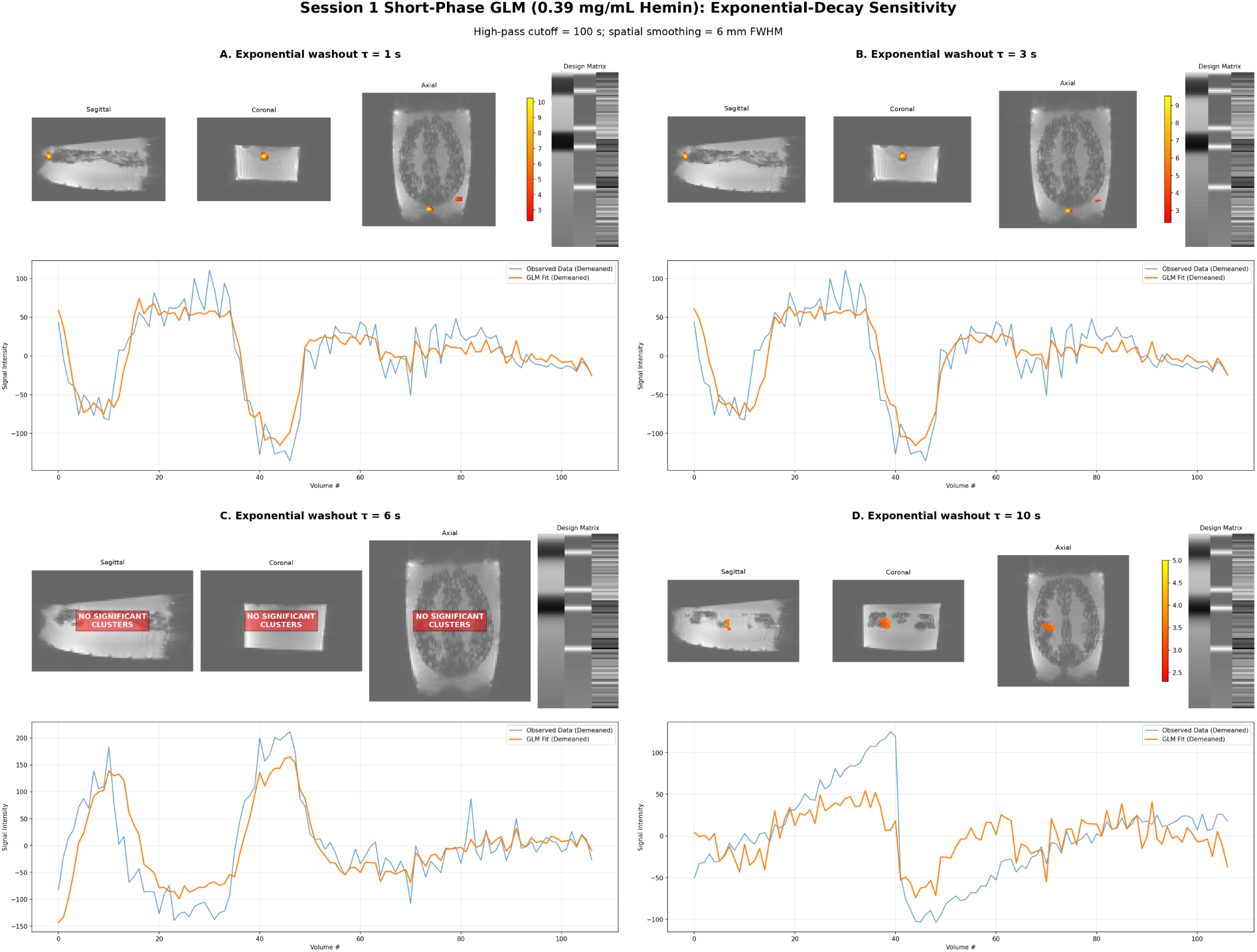
Sensitivity of the Session 1 short-phase 0.39 mg/mL hemin GLM result to the exponential washout time constant. Panels show results obtained using washout time constants of (A) *τ* = 1 s, (B) *τ* = 3 s, (C) *τ* = 6 s, and (D) *τ* = 10 s, while spatial smoothing and the high-pass cutoff were held fixed at 6 mm FWHM and 100 s, respectively. Each panel presents thresholded GLM *Z*-statistic maps in representative sagittal, coronal, and axial views, together with the demeaned observed signal time course (blue) and demeaned GLM fit (orange). Where no cluster survived correction, this is indicated on the anatomical views and the time course was evaluated at the uncorrected statistical peak. All images were derived from the gradient-echo-distortion-corrected functional data.

The detection performance observed here compares favourably with prior active-phantom designs: whereas current-carrying-conductor approaches [19, 20] require dedicated electrical hardware and produce field perturbations whose spatial profile can diverge from distributed in vivo susceptibility patterns, and bubble-compression approaches [21] rely on mechanical actuation with more constrained control over signal timing, the hemin-infusion mechanism used here allows continuous, graded control over contrast-agent concentration and timing using standard clinical infusion hardware. Similarly, unlike the static concentrations used in prior dual-compartment gel designs [22], the present system permits gradual, temporally varying dilution of the paramagnetic agent, more closely approximating the dynamic character of a haemodynamic response, albeit without reproducing its underlying biology. We also avoid the rigid internal partitions used in some prior designs [22, 23], which have been reported to introduce susceptibility artefacts and signal voids at tissue-compartment boundaries not present in vivo.

The central advantage of an anthropomorphic signal-generating phantom over conventional uniform test objects lies in the ability to process the phantom data through the same spatial normalisation pipeline applied to human participants. Because the phantom’s external geometry is derived from an individual’s anatomy, the phantom’s functional data can be coregistered to that participant’s *T*_1_-weighted anatomy using rigid-body transformations (Figure 3) and, from there, optionally mapped into standard stereotaxic space by composing those rigid transforms with the participant’s existing human-to-MNI transforms (Figure 4). This capability is not available with geometrically regular phantoms, whose shape bears no correspondence to any brain template. With the SiGn phantom, one can therefore trace a known input signal, whose spatial extent, timing, and amplitude are controlled by the experimenter, through every stage of a typical fMRI preprocessing and analysis pipeline: motion correction, susceptibility distortion correction, spatial smoothing, temporal filtering, coregistration, and normalisation to MNI space. Any distortion, displacement, or attenuation of the ground-truth signal at the output of the pipeline can be unambiguously attributed to the processing step that introduced it.

This property opens several avenues for methodological investigation. First, it permits systematic evaluation of how different distortion-correction strategies, for example gradient-echo versus spin-echo field-map-based approaches as demonstrated in Session 2 of the present dataset, affect the spatial fidelity of activation maps in a setting where the true activation locus is known. Second, it allows the impact of spatial smoothing kernel width, interpolation scheme, and registration algorithm on effect-size estimates and cluster extent to be quantified against a physical ground truth, rather than inferred from simulations or split-half reliability analyses in human data. Third, standardising the phantom to MNI coordinates enables direct comparison of activation maps acquired on different scanners or after hardware or software upgrades, because any shift in the normalised activation locus would reflect a change in the acquisition or processing chain rather than biological variability between participants.

Beyond its validated functional performance, the primary contribution of this phantom lies in its open, reproducible design, which we discuss below.

The open-data and open-methods philosophy adopted here is integral to these goals. By releasing the STL model files, gel recipes, BIDS-formatted raw data, and version-controlled analysis scripts, we aim to lower the barrier to entry for groups wishing to establish dynamic phantom-based QA programmes. Because the anatomical model is derived from a specific participant whose structural scans are also provided, other laboratories can replicate not only the phantom but the entire registration chain to MNI space, ensuring that cross-site comparisons are performed in a common coordinate frame.

Several limitations of the current work should be acknowledged. First, the signal mechanism, diffusion and flow of a paramagnetic solution through a tube embedded in agar gel, differs fundamentally from the neurovascular coupling that underlies the BOLD effect. The spatial spread of the contrast agent is governed by laminar flow and molecular diffusion rather than by the vascular architecture, and the temporal dynamics are determined by infusion rate and tube geometry rather than by haemodynamic response kinetics. The SiGn phantom is therefore not intended to model the biophysics of the BOLD response; rather, it provides a controlled 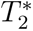 perturbation whose detectability by standard analysis tools can be assessed. A related geometric distinction concerns the physical basis of the 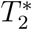 contrast itself: in vivo, BOLD-related dephasing arises around a large population of small, randomly oriented microvessels, whereas in the present phantom the contrast agent is confined to a single, comparatively large-bore tube. This difference in susceptibility geometry means the detailed relationship between signal change and echo time cannot be assumed to mirror the distributed dephasing regime seen in vivo, even though the resulting signal remains a controlled and detectable 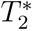 perturbation. Second, the present study examined a single axial slab rather than a whole-brain volume. While extending the design to multiple slabs is straightforward in principle, practical challenges related to gel injection, air bubble formation, infusion-line routing, and print fidelity at whole-brain scale remain to be addressed. Third, the agar-based gels, while stable over the timescale of a scanning session, are subject to gradual dehydration and microbial degradation over days to weeks, limiting the phantom’s useful lifespan. Fourth, the *T*_1_ and flow confounds observed in the infusion tube core indicate that the current single-echo acquisition does not cleanly isolate 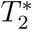 effects; multi-echo sequences or flow-sensitised acquisitions would improve the specificity of future measurements.

Future development of the SiGn phantom could proceed along several directions. Incorpo-ration of multiple, independently addressable infusion channels would permit simultaneous simulation of activations in distinct cortical regions, enabling tests of functional connectivity analyses and network-level statistics. Coupling the infusion system to a programmable syringe pump with event-locked timing would allow convolution of the infusion profile with a canonical haemodynamic response function, producing temporal dynamics that more closely resemble those observed in vivo. Extension to a multi-slab or full-brain geometry, potentially using modular interlocking segments, would increase the anatomical coverage and enable assessment of through-plane registration accuracy. Finally, integration of the phantom into longitudinal scanner monitoring programmes, scanning the device at regular intervals alongside existing ACR or FBIRN protocols, could provide a dynamic complement to the static stability metrics currently in widespread use [15, 16, 39].

The SiGn phantom and its accompanying open pipeline provide a novel tool for fMRI quality assurance that bridges the gap between static test objects and living participants. By embedding a controllable, ground-truth signal within an anatomically realistic geometry that can be spatially normalised to standard brain templates, the phantom enables rigorous, quantitative assessment of the effects of acquisition parameters and preprocessing choices on fMRI activation maps.

## Data and code availability

All data and code supporting this work are openly available on the University of Malta’s Institutional Repository (drUM) at https://doi.org/10.60809/drum.31411158. This includes the BIDS-formatted raw and derivative imaging data, preprocessing and analysis scripts, the containerised reproducibility workflow, 3D-printable STL model files for the anatomical phantom, and tissue-mimicking gel recipes. The reproducibility container is distributed via Docker Hub and archived on Zenodo at https://doi.org/10.5281/zenodo.19495290. All data are shared under the Creative Commons Attribution 4.0 International (CC BY 4.0) license. All code developed by the SARA Project authors for data processing and analysis is released under the BSD 3-Clause license. The archived container additionally bundles third-party software that remains subject to its own upstream license terms; inclusion in the container does not imply relicensing of those components by the authors. Users are responsible for complying with the applicable third-party licenses, including any restrictions that may apply to commercial use of specific bundled software.

## Author contributions

Sephora Galea: writing–original draft; writing–review and editing; building phantom; data analysis.

Brandon Seychell: writing–original draft; writing–review and editing; building phantom; data analysis.

Paola Galdi: writing–review; funding acquisition.

Therese Hunter: building phantom; supervision.

Claude J Bajada: conceptualisation; funding acquisition; writing–original draft; writing– review and editing; data analysis; supervision.

## Acknowledgments

This work is a product of the Synthetic Anatomy for Radiological Applications - Generating a Functional MRI Phantom (SARA) Project, which is part of the Research Excellence Programme by Xjenza Malta. The research team would like to thank the University of Malta’s MRI Platform (UMRI) for access to scanning equipment and services, use of its infrastructure, and resources such as documentation and preprocessing scripts. We also thank the University of Malta’s Data Integrity and Stewardship Cluster (DISC) for support with data management.

## References

[1] Domenico Zacà, et al. “ReStNeuMap: a tool for automatic extraction of resting-state functional MRI networks in neurosurgical practice”. In: Journal of Neurosurgery (2018), pp. 1–8. doi: 10.3171/2018.4.JNS18474.

[2] Donna Dierker et al. “Resting-state functional magnetic resonance imaging in presurgical functional mapping: sensorimotor localization”. In: Neuroimaging Clinics of North America 27.4 (2017), pp. 621–633. doi: 10.1016/j.nic.2017.06.011.

[3] Hanani Abdul Manan, Elizabeth A Franz, and Noorazrul Yahya. “The utilisation of resting-state fMRI as a pre-operative mapping tool in patients with brain tumours in comparison to task-based fMRI and intraoperative mapping: a systematic review”. In: European Journal of Cancer Care 30.4 (2021), e13428. doi: 10.1111/ecc.13428.

[4] Hamad Yahia Abu Mhanna, et al. “Systematic review of functional magnetic resonance imaging (fMRI) applications in the preoperative planning and treatment assessment of brain tumors”. In: Heliyon 11.3 (2025), e42464. doi: 10.1016/j.heliyon.2025.e42464.

[5] Qingbao Yu and Vince D Calhoun. “Resting-state functional network disturbances in schizophrenia”. In: Brain Network Dysfunction in Neuropsychiatric Illness. Ed. by Vaibhav A Diwadkar and Simon B Eickhoff. Cham: Springer, 2021, pp. 217–236. doi: 10.1007/978-3-030-59797-9_10.

[6] Samuele Cortese. “Understanding the network bases of ADHD: an overview of the fMRI evidence”. In: Brain Network Dysfunction in Neuropsychiatric Illness. Ed. by Vaibhav A Diwadkar and Simon B Eickhoff. Cham: Springer, 2021. doi: 10.1007/978-3-030-59797-9_16.

[7] Seiji Ogawa et al. “Brain magnetic resonance imaging with contrast dependent on blood oxygenation”. In: Proceedings of the National Academy of Sciences 87.24 (1990), pp. 9868–9872. doi: 10.1073/pnas.87.24.9868.

[8] Stefano Delli Pizzi et al. “BOLD cardiorespiratory pulsatility in the brain: from noise to signal of interest”. In: Frontiers in Human Neuroscience 17 (2024), p. 1327276. doi: 10.3389/fnhum.2023.1327276.

[9] Paul A Taylor et al. “A set of fMRI quality control tools in AFNI: systematic, in-depth, and interactive QC with afni proc.py and more”. In: Imaging Neuroscience 2 (2024), pp. 1–39. doi: 10.1162/imag_a_00246.

[10] Rebecca J Lepping et al. “Quality control in resting-state fMRI: the benefits of visual inspection”. In: Frontiers in Neuroscience 17 (2023), p. 1076824. doi: 10.3389/fnins.2023.1076824.

[11] Karl F Stupic et al. “A standard system phantom for magnetic resonance imaging”. In: Magnetic Resonance in Medicine 86.3 (2021), pp. 1194–1211. doi: 10.1002/mrm.28779.

[12] Edward F. Jackson et al. Acceptance Testing and Quality Assurance Procedures for Magnetic Resonance Imaging Facilities. AAPM Report 100. American Association of Physicists in Medicine, 2010.

[13] Kathryn E. Keenan et al. “Quantitative magnetic resonance imaging phantoms: A review and the need for a system phantom”. In: Magnetic Resonance in Medicine 79.1 (2018), pp. 48–61. doi: 10.1002/mrm.26982.

[14] Gisella Gennaro et al. “Phantom studies in medical imaging (PSMI): a guide with recommendations and checklist”. In: European Radiology Experimental 9 (2025), p. 105. doi: 10.1186/s41747-025-00641-7.

[15] Lee Friedman and Gary H Glover. “Report on a multicenter fMRI quality assurance protocol”. In: Journal of Magnetic Resonance Imaging 23.6 (2006), pp. 827–839. doi: 10.1002/jmri.20583.

[16] Gary H Glover et al. “Function biomedical informatics research network recommendations for prospective multicenter functional MRI studies”. In: Journal of Magnetic Resonance Imaging 36.1 (2012), pp. 39–54. doi: 10.1002/jmri.23572.

[17] Andreas Altermatt et al. “Design and construction of an innovative brain phantom prototype for MRI”. In: Magnetic Resonance in Medicine 81.2 (2019), pp. 1165–1171. doi: 10.1002/mrm.27464.

[18] Sossena Wood et al. “Design and fabrication of a realistic anthropomorphic heterogeneous head phantom for MR purposes”. In: PLOS ONE 12.8 (2017), e0183168. doi: 10.1371/journal.pone.0183168.

[19] Tiao Chen et al. “BOLD signal simulation and fMRI quality control based on an active phantom: a preliminary study”. In: Medical & Biological Engineering & Computing (2020). doi: 10.1007/s11517-020-02133-9.

[20] Ville Renvall. “Functional magnetic resonance imaging reference phantom”. In: Magnetic Resonance Imaging 27.5 (2009), pp. 701–708. doi: 10.1016/j.mri.2008.11.007.

[21] Akihiro Yamashiro, Takaaki Saito, and Tosiaki Miyati. “Development of a novel task-based functional magnetic resonance imaging phantom based on a bubble-compression approach”. In: Medical Physics (2022). doi: 10.1002/mp.15599.

[22] Johan Olsrud, et al. “A two-compartment gel phantom for optimization and quality assurance in clinical BOLD fMRI”. In: Magnetic Resonance Imaging 26 (2008), pp. 279–286.

[23] Lijun Deng et al. “Anthropomorphic Head MRI Phantoms: Technical Development, Brain Imaging Applications, and Future Prospects”. In: Journal of Magnetic Resonance Imaging (2025). doi: 10.1002/jmri.29818.

[24] Kalina V. Jordanova, Stephen E. Russek, and Kathryn E. Keenan. “Open-source, customizable phantom for low-field magnetic resonance imaging”. In: *Magnetic Resonance Materials in Physics*, Biology and Medicine 38.4 (2025), pp. 727–739. doi: 10.1007/s10334-025-01270-2.

[25] Sephora Galea et al. *The Signal Generating (SiGn) fMRI Phantom: Data, Code, and Fabrication Resources*. University of Malta Institutional Repository (drUM), 2026. doi: 10.60809/drum.31411158.

[26] Sephora Galea et al. The Signal Generating (SiGn) fMRI Phantom: Reproducibility Container. 2026. doi: 10.5281/zenodo.19495290.

[27] Oscar Esteban et al. “fMRIPrep: a robust preprocessing pipeline for functional MRI”. In: Nature Methods 16.1 (2019), pp. 111–116. doi: 10.1038/s41592-018-0235-4.

[28] Kenneth A. Kraft et al. “An MRI phantom material for quantitative relaxometry”. In: Magnetic Resonance in Medicine 5.6 (1987), pp. 555–562. doi: 10.1002/mrm.1910050606.

[29] Vassilios S. Vassiliou, Elaine L. Heng, Peter D. Gatehouse, et al. “Magnetic resonance imaging phantoms for quality-control of myocardial T1 and ECV mapping: specific formulation, long-term stability and variation with heart rate and temperature”. In: Journal of Cardiovascular Magnetic Resonance 18 (2016), p. 62. doi: 10.1186/s12968-016-0275-9.

[30] Marcos Wolf et al. “Reproducible phantom for quality assurance in abdominal MRI focussing kidney imaging”. In: Frontiers in Physics 10 (2022), p. 993241. doi: 10.3389/fphy.2022.993241.

[31] D. C. Grenoble and H. G. Drickamer. “The effect of pressure on the oxidation state of iron. 3. Hemin and hematin”. In: Proceedings of the National Academy of Sciences of the United States of America 61.4 (1968), pp. 1177–1182.

[32] Mark Jenkinson et al. “FSL”. In: NeuroImage 62.2 (2012), pp. 782–790. doi: 10.1016/j.neuroimage.2011.09.015.

[33] Xiangrui Li et al. “The first step for neuroimaging data analysis: DICOM to NIfTI conversion”. In: Journal of Neuroscience Methods 264 (2016), pp. 47–56. doi: 10.1016/j.jneumeth.2016.03.001.

[34] Krzysztof J Gorgolewski et al. “The brain imaging data structure, a format for organizing and describing outputs of neuroimaging experiments”. In: Scientific Data 3 (2016), p. 160044. doi: 10.1038/sdata.2016.44.

[35] Vladimir S. Fonov et al. “Unbiased average age-appropriate atlases for pediatric studies”. In: NeuroImage 54.1 (2011), pp. 313–327. issn: 1053-8119. doi: 10.1016/j.neuroimage.2010.07.033.

[36] Vladimir S. Fonov et al. “Unbiased nonlinear average age-appropriate brain templates from birth to adulthood”. In: NeuroImage 47.Supplement 1 (2009). Organization for Human Brain Mapping 2009 Annual Meeting, S102. doi: 10.1016/S1053-8119(09)70884-5.

[37] Jesper L R Andersson, Stefan Skare, and John Ashburner. “How to correct susceptibility distortions in spin-echo echo-planar images: application to diffusion tensor imaging”. In: NeuroImage 20.2 (2003), pp. 870–888. doi: 10.1016/S1053-8119(03)00336-7.

[38] Mark W Woolrich et al. “Temporal autocorrelation in univariate linear modeling of FMRI data”. In: NeuroImage 14.6 (2001), pp. 1370–1386. doi: 10.1006/nimg.2001. 0931.

[39] David B. Keator et al. “The Function Biomedical Informatics Research Network Data Repository”. In: NeuroImage 124.Pt B (2016), pp. 1074–1079. doi: 10.1016/j.neuroimage.2015.09.003.

